# Estimation of the Hemoglobin Glycation Rate Constant

**DOI:** 10.1101/652818

**Authors:** Masashi Kameyama, Toshika Okumiya, Shinji Tokuhiro, Yoshihisa Matsumura, Hirotaka Matsui, Yasuhiro Ono, Tsuyoshi Iwasaka, Kazuyuki Hiratani, Masafumi Koga

## Abstract

**Aim:** In a previous study, a method of obtaining mean erythrocyte age (*M*_*RBC*_) from HbA1c and average plasma glucose (AG) was proposed. However, the true value of the hemoglobin glycation constant (*k*_*g*_ dL/mg/day), required for this model has yet to be well characterized. Another study also proposed a method of deriving *M*_*RBC*_ from erythrocyte creatine (EC). Utilizing these formulae, this study aimed to determine a more accurate estimate of *k*_*g*_.

**Methods:** 107 subjects including 31 patients with hemolytic anemia and 76 subjects without anemia were included in this study. EC and HbA1c data were analyzed, and *M*_*RBC*_ using HbA1c, AG and the newly-derived constant, *k*_*g*_ were compared to *M*_*RBC*_ using traditional ^51^Cr in three patients whose data were taken from previous case studies.

**Results:** A value of 7.0 × 10^−6^ dL/mg/day was determined for *k*_*g*_. *M*_*RBC*_ using HbA1c, AG and *k*_*g*_ were found to no be significantly different (paired *t*-test, *p* = 0.45) to *M*_*RBC*_ using traditional ^51^Cr.

**Conclusions:** *k*_*g*_ enables the estimation of *M*_*RBC*_ from HbA1c and AG.

## 1 Introduction

HbA1c is widely used as both an indicator of glycemic control, as well as a diagnostic index, for diabetes in clinical settings [1, 2]. Although HbA1c is generally indicative of recent glycemic control over the past 1–2 months, it is known to show reduced correlation to glycemic control status in the presence of diseases which result in a shortened erythrocyte lifespan such as hemolytic anemia [3].

We have recently proposed a simple method to obtain mean erythrocyte age (*M*_*RBC*_) from HbA1c and average glucose (AG) [4]:

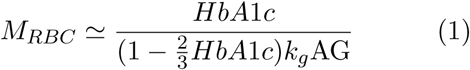

where *k*_*g*_ is the rate constant of the glycation reaction. This formula provides meaningful information for the diagnosis of anemia. We estimated *k*_*g*_ to be 6–10×10^−6^ dL/mg/day based on past literature [4]. However, a more accurately estimated value of *k*_*g*_ would provide more useful information.

The relationship between *M*_*RBC*_ and erythrocyte creatine (EC) was previously established based on a model [5] as following:

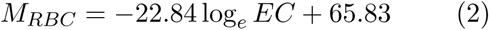

This study aimed to determine the accurate value of *k*_*g*_ from EC-derived *M*_*RBC*_ and HbA1c.

## 2 Materials and Methods

### 2.1 Participants

107 subjects including 31 patients with hemolytic anemia and 76 subjects without anemia were included in this study. All samples were prepared and analyzed in accordance with the protocols approved by the institutional committees at Kumamoto University and other collaborating institutions.

Patients with hemolytic anemia were recruited from 115 patients who were older than 20 years old and required laboratory tests including complete blood counts and reticulocyte counts (ret) for clinical reasons. Those who were suspected of having diabetes mellitus (DM) based on history, a low 1,5-AG value (male, *<*14.9 *µ*g/mL; female, *<*12.4 *µ*g/mL), or had comorbid liver or renal diseases, were excluded, as liver and renal diseases affect HbA1c and glycated albumin (GA). EC, HbA1c, GA, haptoglobin, and other biochemical screening items were measured using the existing plasma samples from these patients. Use of existing plasma samples from anemic patients without written consent was approved by the institutional review board.

Participants without anemia were recruited from medical examination checkup recipients at Takagi Hospital. Those who had anemia, DM, liver disease, renal disease or who were pregnant were excluded to avoid confounding effects on HbA1c or GA value. We provided the healthy volunteers with detailed information about the study, and all participants without anemia provided written informed consent to participate.

### 2.2 Data interpretation

HbA1c expressed in International Federation of Clinical Chemistry (IFCC) units (iA1c) was used for calculations in this study. While the National Glycohemoglobin Standardization Program (NGSP) is used to express HbA1c in many clinical research and medical care settings, NGSP is measured by an old standardized method and at the time of conception, high performance liquid chromatography (HPLC) was not able to distinguish true HbA1c from other products. HPLC technology later advanced, however the derived HbA1c value is adjusted to NGSP in the interest of consistency. IFCC provides a strict definition of iA1c as hemoglobin with a glycated valine in the *β*-chain. Thus, iA1c value is preferred value for estimation of erythrocyte glycation.

To acquire iA1c from HbA1c expressed in NSGP unit, we used the following equation [6]:

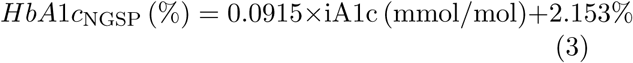

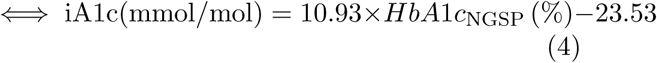

*M*_*RBC*_ was acquired from EC by the aforementioned equation (2).

An AG value of 100 mg/dL was substituted for blood glucose values derived using CGM. This number was based on the average AG of non-diabetic participants and the previously reported findings from a study which showed the median AG in healthy subjects to be reported to be 101.0 (96.3 – 106.0) mg/dL [7] and another ADAG (A1c-Derived Average Glucose) study which found that the AG of the non-diabetic group of their study was similarly 100mg/dL [8, 9].

*M*_*RBC*_ was also determined using ^51^Cr half-life. As the reference range for ^51^Cr half-life was described as 28–30 days [10], 30±5 days [11], and 26–40 days [12], *M*_*RBC*_ was calculated by multiplying ^51^Cr half-life and 2.14 (=60/28), 60 days being the normal value for *M*_*RBC*_.

### 2.3 Data analysis

*EC* and *M*_*RBC*_ data were analyzed using a spreadsheet software, Excel^®^ 365 (Microsoft Corporation, Redmond, WA, USA).

### 2.4 Estimation of *k*_*g*_

The following two methods were used to estimate *k*_*g*_. The slope method – the following equation (5) derived from equation (1) shows that the slope of the line connecting a point and the origin is *k*_*g*_AG.

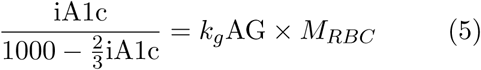

Estimating the slope of the regression line through the origin by the least square model:

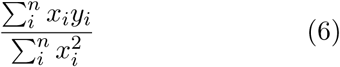

where *x*_*i*_, *y*_*i*_ are *M*_*RBC*_ and 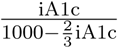 of each participant, respectively.

The direct method – the *k*_*g*_ of each participant was calculated by the following equation:

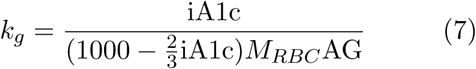

Then, average and standard deviation of each *k*_*g*_ was calculated.

### 2.5 Confirmation of derived *k*_*g*_

The method of obtaining *M*_*RBC*_ from *AG* and iA1c was applied to data from three patients with latent hemolysis who were presented in a previous case studies [10–12].

Herranz [10] and Ishii’s [11] data showed changes in HbA1c during the course of the study. Therefore, *M*_*RBC*_ was calculated separately for each period. For the Ishii case [11], AG was calculated by averaging self-monitoring of blood glucose (SMBG) data for each periodperiod. The Hiratani study [12] examined ^51^Cr erythrocyte lifespan measurement during hospitalization in Oct 1999 and CGM in Feb 2016. While HbA1c and plasma glucose concentrations fluctuate routinely, RBC lifespan remain comparatively constant, especially when influenced by a certain diseases (stomatocytosis). Furthermore, supply of ^51^Cr was ceased in Japan in 2015 and thus it can no longer by used to study erythrocyte lifespan.

## 3 Results

### 3.1 Participant characteristics

Participant demographics are shown in Table 1. All participants had no more than 16% GA. There was no significant difference in the GA of anemic and non-anemic subjects. However HbA1c, Hb, EC and their derivatives showed significant variation between the two groups.

**Table 1:** Participants Characteristics.

| | non-hemolysis | hemolysis | $p$ |
| --- | --- | --- | --- |
| $n$ (M/F) | 76 (30/46) | 31 (17/14) | 0.1463 |
| Age (years) | 62.3±7.9 | 45.6±15.0 | $1.37 \times 10^{-6}$ |
| HbA1c (%) | 5.78 ± 0.25 | 4.05 ± 0.78 | $2.27 \times 10^{-13}$ |
| iA1c (mmol/mol) | 39.7 ± 2.7 | 20.8 ± 8.5 | $2.27 \times 10^{-13}$ |
| GA (%) | 13.57 ± 1.07 | 13.06 ± 1.75 | 0.151 |
| GA/iA1c | 0.343 ± 0.032 | 0.762 ± 0.363 | $5.85 \times 10^{-7}$ |
| Hb (g/dL) | 14.26 ± 1.16 | 9.75 ± 1.97 | $2.44 \times 10^{-14}$ |
| EC (μmol/g Hb) | 1.40 ± 0.21 | 5.47 ± 2.13 | $1.42 \times 10^{-11}$ |
| EC-$M_{RBC}$ (days) | 58.5 ± 3.41 | 29.0 ± 10.0 | $4.46 \times 10^{-17}$ |

Results are expressed as mean ± standard deviation (SD). Sex ratio was examined by $\chi^2$ test. Other items were examined by $t$ -test (bilateral). GA, glycated albumin; EC, erythrocyte creatine

The demographic information on the 3 patients from the previous cases are shown in Table 2.

**Table 2:** Characteristics of three reported patients with latent hemolysis and DM.

| Case | Herranz [10] | Ishii [11] | Hiratani [12] |
| --- | --- | --- | --- |
| Age/Sex | 30F | 72M | 58F |
| Disease | AIHA | AIHA | HSt |
| DM | Type 1 | Type 2 | Type 2 |
| HbA1c (%) | 5.4 | 6.5 | 5.8 |
| GA (%) | — | 26.1 | 23.3 |
| Hb (g/dL) | normal | 13.5 | 11.5 |
| Ret (%) | normal | 1.3 | 1.3 |
| Hpt (mg/dL) | normal | 82 | 58 |

These patients showed normal Hb, reticulocyte, and haptoglobin. AIHA, autoimmune hemolytic anemia; HSt, hereditary stomatocytosis; DM, diabetes mellitus; GA, glycated albumin; Hb, hemoglobin; Ret, reticulocyte; Hpt, haptoglobin.

### 3.2 Estimation of *k*_*g*_

EC derived *M*_*RBC*_ and 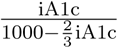 are shown in Figure 1. A linear relationship was successfully observed.

**Figure 1.**
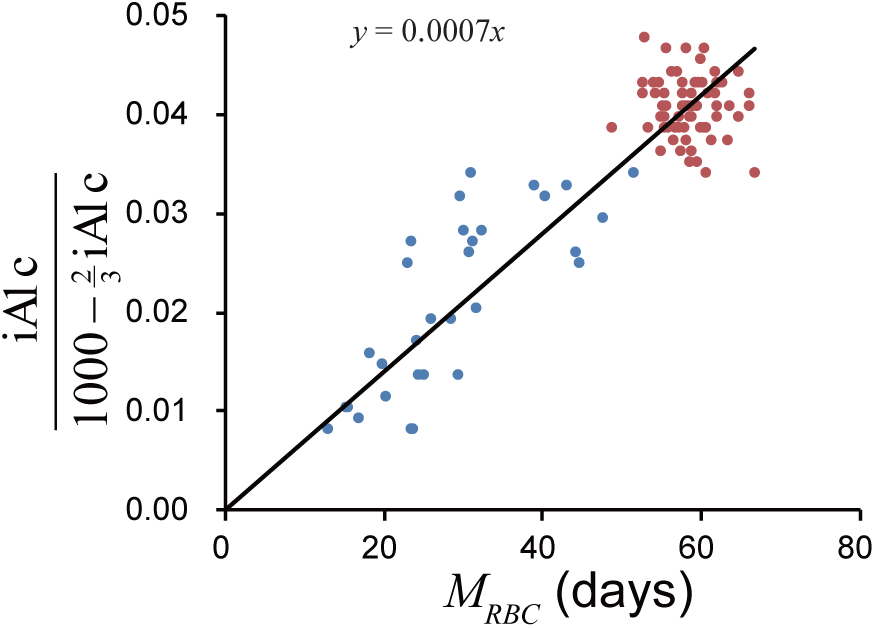
Relationship between EC derived *M*_*RBC*_ and iA1c/(1000−(2/3)iA1c). *Red circles* denote non-hemolytic participants and *blue circles* denote hemolytic patients. *Black line* denotes regression line through origin. (For interpretation of the references to color in this figure legend, the reader is referred to the web version of this article.)

*k*_*g*_ calculated by the two methods outlined previously, for non-hemolytic participants and the entire study population are seen in Table 3. All 4 figures can be approximated to 7 × 10^−6^. Figure 1 shows that data from severe hemolytic patients is less stable. Thus, the value derived from the direct method for calculating *k*_*g*_ is likely to be the least accurate. Excluding this value as an outlier, the 3 remaining figures were 6.94 – 6.99 ×10^−6^ (average 6.970 × 10^−6^). Therefore, considering significant figures, *k*_*g*_ can be said to be 7.0 × 10^−6^.

**Table 3:** *k*_*g*_ estimation

| population | slope | straight forward |
| --- | --- | --- |
| the whole | $6.973 \times 10^{-6}$ | $(7.073 \pm 1.229) \times 10^{-6}$ |
| non-hemolytic | $6.942 \times 10^{-6}$ | $(6.994 \pm 0.662) \times 10^{-6}$ |

Results obtained by the direct method are expressed as mean ± standard deviation (SD).

### 3.3 Confirmation of derived *k*_*g*_

The *M*_*RBC*_ using the derived *k*_*g*_, 7.0 × 10^−6^ and *M*_*RBC*_ using ^51^Cr half-life are shown in Table 4.

**Table 4:** *M*_*RBC*_ of 3 cases in literature

| Case | Herranz [10] |  | Ishii [11] |  | Hiratani [12] |
| --- | --- | --- | --- | --- | --- |
| HbA1c (%) | 5.8 | 4.9 | 6.0 | 6.7 | 6.5 |
| iA1c (mmol/mol) | 39.9 | 30.0 | 42.3 | 50.2 | 47.5 |
| AG (mg/dL) | 203 | 148 | 138 | 177 | 184 |
| iA1c derived $M_{RBC}$ | 28.7 | 29.6 | 45.3 | 41.9 | 38.1 |
| (days) | 29.2 |  | 43.6 |  | 38.1 |
| $^{51}\text{Cr}$ half-life (days) | 20 | | 20 | | 17.8 |
| $^{51}\text{Cr}$ derived $M_{RBC}$ (days) | 42.9 | | 42.9 | | 38.1 |

*M*_*RBC*_ derived from iA1c were 36.95±5.93, *M*_*RBC*_ derived from ^51^Cr half-life were 41.29±2.22. Paired *t*-test: *t*, −0.9278; df, 2; *p* (bilateral), 0.4514. Thus, *MRBC* derived from iA1c and *M*_*RBC*_ using ^51^Cr half-life were not significantly different.

## 4 Discussion

Based on EC-derived *M*_*RBC*_ and HbA1c data, a more accurate value for the constant *k*_*g*_ was obtained. Though *k*_*g*_ was previously determined to be 6–10×10^−6^ dL/mg/day [4], the more accurate value of 7.0 × 10^−6^ improves the usefulness of the proposed model allowing closer approximation of *M*_*RBC*_ based on AG and iA1c.

Moreover, the validity of *k*_*g*_ has been confirmed through comparison of *M*_*RBC*_ derived from iA1c and *k*_*g*_ with *M*_*RBC*_ derived from ^51^Cr half-life. Of the three patients with hemolytic anemia and comorbid DM analyzed, data from two patients showed a remarkable correlation with the model derived figures. Data from one patient showed a 1.47 times difference in values however, this may be attributable to the use of SMBG instead of CGM, and the difficulty of standardizing 51Cr data containing elution.

Variant hemoglobin should be distinguished from hemolysis when *M*_*RBC*_ determined by equation (1) is low. Glycated variant hemoglobin will exhibit different peaks in HPLC from normal HbA1c, resulting in erroneously low values for HbA1c (some variants show an artefactually high value). It has previously been reported that variant hemoglobin can be detected by the dissociation between HbA1c measured by HPLC and by immunoassay [13]. Moreover, some variant hemoglobins such as Hb Himeji [14] have different *k*_*g*_ values from normal Hb. In patients with these variant hemoglobins, equation (1) is likely to provide a falsely low *M*_*RBC*_.

There are a number of limitations to this study. The data used to calculate a more specific estimate of *k*_*g*_ contained EC and HbA1c, but lacked CGM data, necessitating the use of. 100 mg/dL as an approximation of AG. However, participants were confirmed to be free of DM through GA, an indicator of glycemic control that is independent of mean erythrocyte age, with a cut off of GA no more than 16%. Further study with more complete data including CGM, HbA1c and EC would provide an even more definitive value for *k*_*g*_. Another limitation is that the value for *k*_*g*_ derived in this study is totally dependent on equation (2) that derives *M*_*RBC*_ from EC. This equation was based on old published data [15], which used less sensitive and poorly specific chemical methods of measuring creatine which were prone to cross-reactivity with other guanidino compounds. This may reduce the reliability of the system. In contrast, in this study creatine was measured using an enzymatic method which was sensitive and specific to creatine in erythrocytes which uses 10-*N*-methylcarbamoyl-3,7-*bis* (dimethylamino) phenothiazine (MCDP), an *N*-methylcarbamoyl derivative of methylene blue, with a high molar absorption coefficient (9.6 × 10^7^L mol^−1^ cm^−1^) [16], as a chromogen.

## Acknowledgement

This study is partly supported by Grants-in-Aid for Scientific Research, 18K07488 for M.Kameyama. The authors would like to thank Ms. Natalie Okawa for English language editing of this manuscript.

## References

[1] Koenig RJ, Peterson CM, Jones RL, Saudek C, Lehrman M, Cerami A. Correlation of glucose regulation and hemoglobin A1c in diabetes mellitus. N Engl J Med. 1976;295:417–420. doi:10.1056/NEJM197608192950804.

[2] American Diabetes Association. Glycemic targets in *Standards of Medical Care in Diabetes – 2017*. Diabetes Care. 2017;40:S48–S56. doi:10.2337/dc17-S009.

[3] Panzer S, Kronik G, Lechner K, Bettelheim P, Neumann E, Dudczak R. Glycosylated hemoglobins (GHb): an index of red cell survival. Blood. 1982;59:1348–1350.

[4] Kameyama M, Takeuchi S, Ishii S. Steadystate relationship between average glucose, HbA1c and RBC lifespan. J Theor Biol. 2018;447:111–117. doi:10.1016/j.jtbi.2018.03.023.

[5] Kameyama M, Koga M, Okumiya T. A novel method for calculating mean erythrocyte age using erythrocyte creatine. bioRxiv. 2019; 642942. doi:10.1101/642942.

[6] Hoelzel W, Weykamp C, Jeppsson JO, et al. IFCC reference system for measurement of hemoglobin A1c in human blood and the national standardization schemes in the United States, Japan, and Sweden: a method-comparison study. Clin Chem. 2004;50:166–174. doi:10.1373/clinchem.2003.024802.

[7] Tsujino D, Nishimura R, Taki K, Miyashita Y, Morimoto A, Tajima N. Daily glucose profiles in Japanese people with normal glucose tolerance as assessed by continuous glucose monitoring. Diabetes Technol Ther. 2009; 11:457–460. doi:10.1089/dia.2008.0083.

[8] Malka R, Nathan DM, Higgins JM. Mechanistic modeling of hemoglobin glycation and red blood cell kinetics enables personalized diabetes monitoring. Sci Transl Med. 2016;8:359ra130. doi:10.1126/scitranslmed.aaf9304.

[9] Nathan DM, Kuenen J, Borg R, Zheng H, Schoenfeld D, Heine RJ. Translating the A1C assay into estimated average glucose values. Diabetes Care. 2008;31:1473–1478. doi:10.2337/dc08-0545.

[10] Herranz L, Grande C, Janez M, Pallardo F. Red blood cell autoantibodies with a shortened erythrocyte life span as a cause of lack of relation between glycosylated hemoglobin and mean blood glucose levels in a woman with type 1 diabetes. Diabetes Care. 1999;22:2085–2086. doi:10.2337/diacare.22.12.2085.

[11] Ishii C, Tane N, Negishi K, Katayama S. A case of type 2 diabetes who showed discrepancy between plasma glucose and HbA1c due to latent autoimmune hemolytic anemia (in Japanese). J Japan Diab Soc. 2001; 44:157–160. doi:10.11213/tonyobyo1958.44.157.

[12] Hiratani K, Natazuka T, Suemori S, Wada H, Koga M. A case of stomatocytosis in a type 2 diabetic patient accompanied with falsely low HbA1c levels due to latent hemolysis (in Japanese). J Japan Diab Soc. 2016;59:719–723. doi:10.11213/tonyobyo.59.719.

[13] Miyazaki A, Kohzuma T, Kasayama S, Koga M. Classification of variant forms of haemoglobin according to the ratio of glycated haemoglobin to glycated albumin. Ann Clin Biochem. 2012;49:441–444. doi:10.1258/acb.2012.011192.

[14] Koga M, Inada S, Shimizu S, Hatazaki M, Umayahara Y, Nishihara E. Aldimine formation reaction, the first step of the maillard early-phase reaction, might be enhanced in variant hemoglobin, Hb Himeji. Ann Clin Lab Sci. 2015;45:643–649.

[15] Fehr J, Knob M. Comparison of red cell creatine level and reticulocyte count in appraising the severity of hemolytic processes. Blood. 1979;53:966–976.

[16] Okumiya T, Jiao Y, Saibara T, et al. Sensitive enzymatic assay for erythrocyte creatine with production of methylene blue. Clin Chem. 1998;44:1489–1496.

